# Conformational drugging pockets at the surface of c-MYC proto-oncongenes

**DOI:** 10.64898/2026.05.28.728384

**Authors:** Yanhong Ge, Huixia Lu, Jordi Marti

## Abstract

The MYC proto-oncogene encodes a master transcription factor essential for regulating cellular homeostasis, growth, and metabolism. However, its deregulation is involved in up to 70% of human cancers, where it drives uncontrolled cellular proliferation. Structurally, MYC is characterized by an N-terminal transactivation domain and a C-terminal bHLHLZ domain critical for DNA binding and dimerization with MAX. Despite its clinical significance, MYC has remained historically ”undruggable” due to its intrinsically disordered structure and the absence of traditional binding pockets. In this study, we combined all-atom Molecular Dynamics and well-tempered Metadynamics simulations to map the conformational landscape of a MYC motif in aqueous ionic solution. The selected energy-minimised structures exhibit stable conformations and drug pockets, providing structural foundations for targetting c-MYC, which is traditionally considered ”undruggable”. Furthermore, both the electrostatic potential and docking results for the obtained structure and the MYC inhibitor 10074-G5 demonstrated excellent fit, highlighting its potential as a drug target. Remarkably, MYC maintained steady binding to 10074-G5 throughout the 500 ns simulation. These findings provide a previously uncharacterized structural foundation for the design of innovative therapeutic interventions, such as small-molecule inhibitors or peptides aimed at disrupting oncogenic MYC activity.

## TOC Graphic

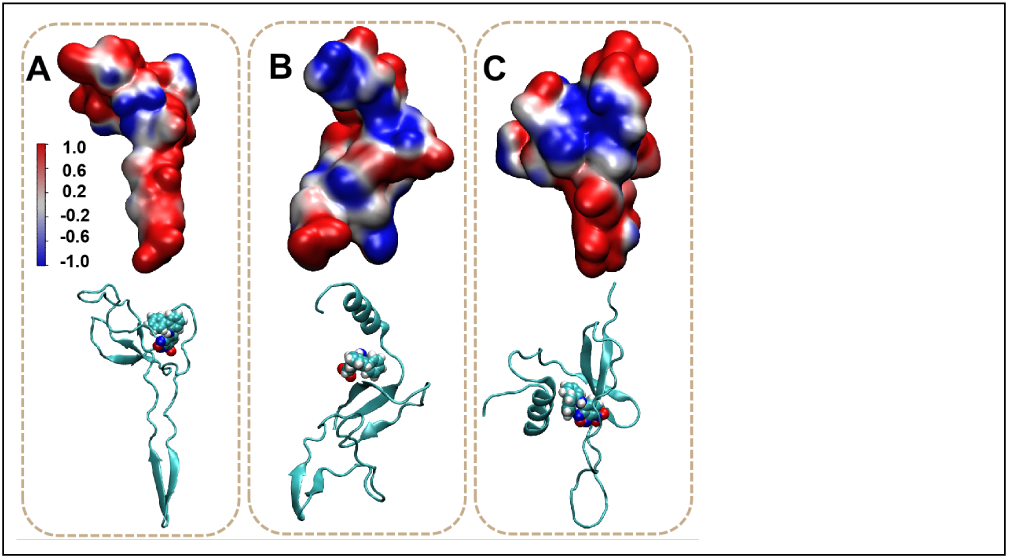

## Introduction

MYC proto-oncogenes encode a transcription factor that plays a crucial role in regulating numerous cellular processes, including growth, differentiation, metabolism, apoptosis, and DNA repair.^1–4^ MYC is vital for normal cellular development, but its overexpression or mutation can drive tumorigenesis. MYC is highly conserved between species,^5^ reflecting its essential role in maintaining cellular homeostasis and cell cycle progression. First identified within the avian retrovirus MC29,^6,7^ the c-MYC gene was one of the first proto-oncogenes to be recognized, and its evolutionary conservation further underscores its biological importance. MYC proteins consist of three key regions: a transactivation domain at the N-terminus, a central region, and the bHLHLZ domain at the C-terminus. The bHLHLZ domain is characteristic of a superfamily of transcription regulators that mediate cellular responses to extracellular and intracellular signals by modulating gene expression.^8^ This domain enables MYC to bind to DNA through E-boxes, specific sequences that facilitate its interaction with various cofactors, particularly the MAX protein.^9,10^

In many cancers, MYC is overexpressed, disrupting the balance between MYC activation and repression. This imbalance favors MYC–MAX complexes, which promote uncontrolled cell growth and inhibit differentiation. Although MYC was once thought to require MAX for both activation and repression, recent studies have shown that MYC retains significant biological activity even in the absence of MAX,^11^further emphasizing the importance of the MYC monomer as a target. MYC expression is deregulated in up to 70% of human cancers,^1^ including Burkitt lymphoma, colorectal cancer, melanoma, prostate cancer, breast cancer (via translocation or amplification of c-MYC) and neuroblastoma (via amplification of n-MYC), among others.^7,12^

Despite its critical role in oncogenesis, targeting MYC directly remains challenging due to its lack of easily druggable sites, since MYC is a disordered protein that does not present obvious binding pockets for small molecule inhibitors. ^13–16^ As a result, strategies to inhibit MYC function have focused on indirect approaches,^1,17,18^ such as disrupting the dimerization of MYC-MAX, preventing MYC-DNA binding, or interfering with MYC target gene expression.^19^ Although direct MYC inhibitors have not yet been approved research continues, with promising therapeutic agents currently under development and undergoing clinical trials. An innovative approach involves Omomyc,^20^ a mutant of MYC engineered to disrupt MYC dimerization. Omomyc contains a modified bHLHLZ domain that allows it to homodimerize and heterodimerize with MYC and MAX, disrupting the normal MYC function. Early clinical trials with Omomyc have shown promising results, indicating that it could be a potential therapeutic option.

In order to locate potential drugging pockets, it is crucial to determine the most stable configurations of MYC, since prototype drugs need to target structures able to be bound and hold long-term durability. In this study, we leverage Molecular Dynamics (MD) simulations at the all-atom level, which provide high temporal resolution and precise force fields to model the behavior of MYC and its interactions with other proteins and drugs. Interestingly, MD can model hydrogens at the classical^21,22^ or quantum levels^23^ although in biological simulations a classical rigid model for water (TIP3P) is usually taken. On the one hand, by analyzing the conformational dynamics of MYC we aim to uncover potential druggable sites and better understand the mechanisms of MYC dimerization and DNA binding. However, the description of the stable states on the free energy hypersurface of isolated MYC in solution, which can be accurately reproduced using tools such as well-tempered metadynamics (WTM) simulations, is a key information to combine with MD data and help the location of pockets or regions at the MYC surface to be targeted, paving the way for future therapeutic interventions based on the design of new drugs. In the present work, structural and free energy data have been combined in order to gather key information on: (1) stable dynamical structures of MYC in aqueous ionic solution, obtained from both MD and WTM, and (2) drugging pockets located on the surface of MYC, which can potentially be targeted by a variety of compounds, some of them being currently designed and evaluated by *in silico* techniques. To validate the stability of the drug pocket, we docked the lowest-energy conformation identified in the WTD with a MYC inhibitor and simulated the resulting complex, thereby providing a structural basis for the design of small-molecule inhibitors.

## Methods

### System preparation and setups

In the present work, we conducted MD and WTM simulations of a motif species c-MYC in NaCl aqueous solution, with the sequence^24^ expressed by 82 amino acids: NVKRRTHNVLER-QRRNELKRSFFALRDQIPELENNEKAPKVVILKKATAYILSVQAEEQKLISEEDLLRKR- REQLKHKLEQL. The selected motif of MYC comprises the specific sequence of MYC bound to MAX in the three following reported crystal structures (CS), downloaded from RCSB PDB Protein Data Bank,^25,26^ namely file names ’1NKP’, ’6G6K’ and ’8OTS’. We simulated six classical MD systems (three long 3 *µ*s simulations and three replicas of length 2 *µ*s for the sake of control). Every system contained one motif MYC protein fully solvated by TIP3P water molecules^27^ in sodium chloride at the human body concentration (0.15 M). The total number of particles in these 6 MD systems ranged between 223305 and 227872 atoms and the corresponding sizes of the simulations boxes were between 133.8×133.6×133.4 °A³ and 135.5×134.9×134.9 °A³, in order to get the MYC motif fully hydrated and located far enough from its replica images. All inputs were generated by means of the CHARMM-GUI solution builder^28–30^ assuming the CHARMM36m force field.^31^ All bonds involving hydrogens were set to fixed lengths, allowing fluctuations of bond distances and angles for the remaining atoms. The MYC proteins in their solvated water box were energy minimized and well equilibrated (NVT ensemble) before generating MD (NPT ensemble) and WTM simulations. MD trajectories were used to feed the initial configurations of the WTM runs. Once we have systematically verified the physical equivalence of the three structures, we have obtained averaged properties in all cases.

The GROMACS/2023 package^32^ was employed for the system minimization as well as for the equilibration and production steps of the MD simulations. The system was minimized with steepest descents and a conjugate-gradient step every 10 steps until convergence to Fmax ≤ 1000 kJ/mol. After energy minimization, a time step of 2 fs was used in all equilibration simulations and the particle mesh Ewald method with Coulomb radius of 1.2 nm was employed to compute long-ranged electrostatic interactions. The cutoff for Lennard-Jones interactions was set to 1.2 nm. The system was equilibrated for 1.25 *µ*s with the Nośe-Hoover thermostat set at 310.15 K in the NVT ensemble, before switching to the NPT ensemble at pressure 1 atm to collect statistics. Periodic boundary conditions (PBC) in the three directions of space were taken in all cases.

### Well-tempered metadynamics

A broad range of methodologies have been proposed to address the challenging task of computing free energy landscapes in multidimensional systems.^33–37^ These approaches aim to capture the intricate behavior of complex systems across multiple degrees of freedom. In the present study, we have utilized the WTM method, which has proven to be particularly effective in exploring free energy surfaces in systems with multiple reaction coordinates. This method has demonstrated significant success in tackling the free energy landscapes of a wide array of complex systems.^38–43^ One of the key advantages of WTM is its versatility, as it can be applied to a broad spectrum of systems, from small molecular systems to larger, more intricate ones. Notably, WTM has shown remarkable efficacy when used to study model cell membrane systems, including those with attached small molecules and proteins. ^44–46^ This makes it particularly well-suited to the investigation of biological processes at a molecular level, which has been a primary focus of recent work in our research group. The ability of WTM to handle the complexity of such systems, including their dynamic interactions and high-dimensional configuration spaces, positions it as a powerful tool for advancing our understanding of molecular mechanisms in various biological contexts.

Extensive WTM simulations considering different collective variables (CV) with total lengths between 7.8 and 10 *µ*s were performed, and once two suitable CV were determined (see section ”Collective Variables” for the choice of CV), we obtained the Gibbs free energy (hyper)surface (FES), starting from the last configuration of one MD simulation (labeled as ”replica 1”, see below), considering a Nośe-Hoover thermostat at 310.15 K and a Parrinello-Rahman barostat at 1 atm in the NPT ensemble. These WTM simulations were performed using the joint GROMACS/2023-plumed-2.9.2 tool. ^47,48^ As in the MD simulations, PBC in the three directions of space were also considered. The total number of atoms in the setups for the WTM simulations was of 39489 with a box size of 75.0×75.0×75.0 °A³.

### Collective Variables

The enhanced sampling method WTM applies a time-dependent biasing potential along a set of CV by adding a Gaussian additional bias potential to the total potential in order to overcome barriers larger than *k_B_T* , with *k_B_* being Boltzmann’s constant and *T* the temperature. In the method, sampling is performed on a selected number of degrees of freedom (*s_i_*), i.e. a chosen set of CV. For each degree of freedom, the biased potential *V* (*s_i_, t*) is a dynamical function constructed as the sum of Gaussian functions:

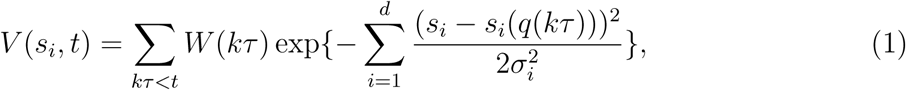

where *k* is an integer, *τ* is the Gaussian deposition stride, *W* (*kτ* ) is the height of the Gaussian and *σ_i_* is the width of the Gaussian for the i-th CV. Further details are provided elsewhere.^49–51^ Unlike unbiased MD simulations, able to track the dynamical evolution of the system around equilibrium states (stable states), the biased potential can force the system to move around all possible states inside a particular range of the subspace of selected CV.

We should point out that determining and using three or more CVs in a WTM simulation is truly unsuitable from the computational point of view. The usual choice is to consider one or two CVs.^52^ The use of two CVs produces a complete enough description of the FES and we have observed this is the optimal way to proceed.^53^ Later on, we will project the free energies onto one single coordinate, integrating out the contribution of the second CV. The values of the parameters taken for the WTM simulations^50^ are listed in Table 1.

**Table 1:** Two-dimensional WTM simulation parameters of aqueous c-MYC systems, where the CVs are *d*1 and *d*2.

| Parameter | Value |
| --- | --- |
| Gaussian width of $d1$ [nm] | 0.082 |
| Gaussian width of $d2$ [nm] | 0.035 |
| Gaussian hill [kJ/mol] | 1.0 |
| Deposition stride [ps] | 20 |
| Biasfactor | 10 |
| Simulation time [μs] | 7.8 |

In this work, we have taken a specific approach similar to previous works ^44,54,55^ where CVs were defined to obtain clear and well defined free energy landscapes. After a detailed study of several distances and angular coordinates, the two CV selected to perform the 2D WTM calculations are shown in Fig. 1. We have selected: (1) as CV1, the distance (*d1* ) between the residue ASN907 within the alpha-helix close to the C-terminal of MYC and the residue HIS976 within the second alpha-helix, close to the N-terminal of MYC; and (2) as CV2 the distance (*d1* ) between residues LEU951 and LEU966, both located at the central part of the second alpha-helix of MYC, in order to study the potential torsions of this particular alpha-helix.

**Figure 1:**
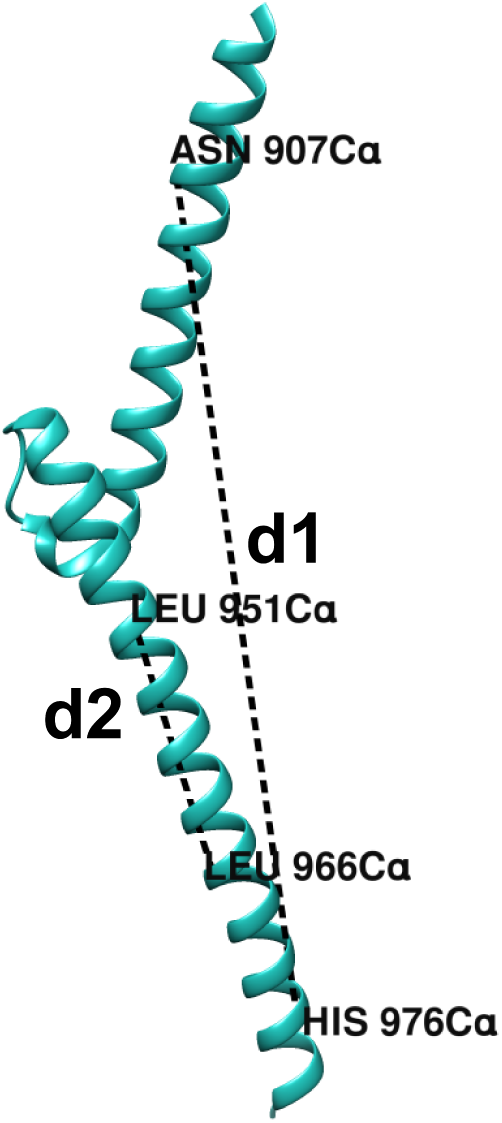
Schematic diagram of the atoms involved in the definition of CVs: (1) the distance (*d1* ) between the ASN907 alpha carbon atom and the HIS976 alpha carbon atom and (2) the distance (*d2* ) between the LEU951 alpha carbon atom and the LEU966 alpha carbon atom.

Specifically, *d*1 is a global variable, while *d*2 serves as a local variable for the long helix region. Since clustering analysis indicates that the ends of the helices tend to uncoil and to avoid interference introduced by the highly flexible terminal residues and the intermediate loop, both d1 and d2 exclude the N-terminus (residues 900–906) and C-terminus (residues 977–981) and loop (residues 927–937), which lack a well-defined structure.

As a relevant remark, we should inform the reader about the importance of setting the correct parameters and specifications for the WTM simulations within the PLUMED project tool employed in the present work. To ensure physically meaningful distance-based CVs during metadynamics simulations, careful treatment of periodic boundary conditions is essential. In PLUMED, distances between atoms or molecular groups may otherwise be evaluated using the nearest periodic image, which can produce artificially short distances when the system spans across the simulation box. In the present work, this issue was resolved by combining the ”WHOLEMOLECULES” directive with the ”NOPBC” option in the CV’s definition. The ”WHOLEMOLECULES” command reconstructs fragmented molecular coordinates across periodic boundaries prior to CV evaluation, while ”NOPBC” ensures that the distance is computed using the actual molecular geometry rather than the minimum-image convention.

Figure 2 illustrates the importance of this treatment. In panel A, where ”WHOLEMOLECULES” and ”NOPBC” were properly applied, the trajectory-derived distance is in excellent agreement with the values reported in the ”COLVAR” file (which stores all values of the 2 CVs during the simulation). This confirms that the CV accurately represents the physical separation between the selected groups. In contrast, panel B shows the result obtained without proper PBC handling, where commands ”WHOLEMOLECULES” and ”NOPBC” were not considered. Further details can be seen in the corresponding Github repository (https://github.com/yanhong77634/myc). In panel B, although the trajectory indicates a progressive increase in the intermolecular distance, the corresponding ”COLVAR” values remain artificially low because PLUMED evaluates the distance to a periodic image of the molecule. These results emphasize that reconstruction of molecular continuity together with explicit disabling of periodic-image distance evaluation is critical for reliable enhanced-sampling simulations involving large conformational displacements or intermolecular separation processes.

**Figure 2:**
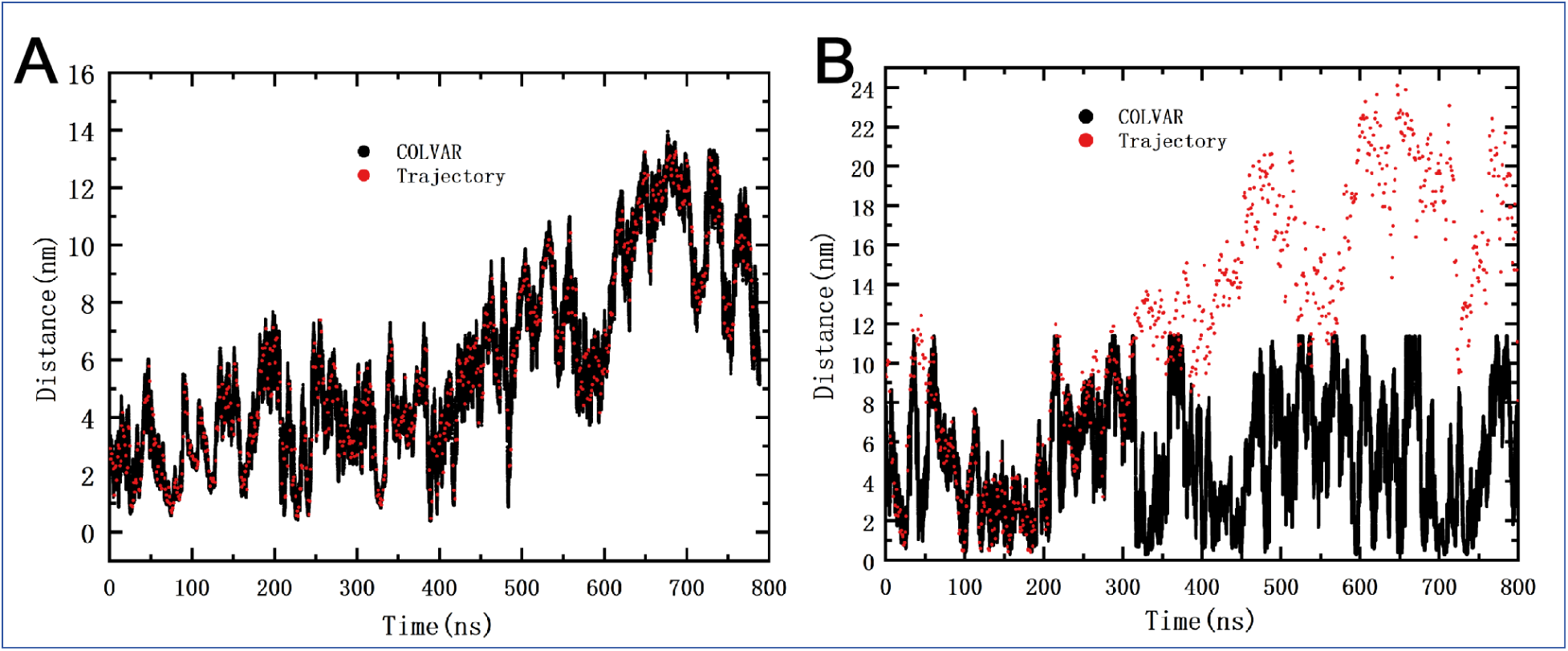
Comparison of values of CV *d*1, that corresponds to the distance between the ASN907 and the HIS976 alpha carbons of MYC in a MD trajectory (red dots) and the values in the ”COLVAR” file calculated by PLUMED (full line) with the correctly configured PLUMED file (A) and using a wrong configuration (B).

### Data analysis

#### Principal Component Analysis

Principal Component Analysis (PCA) is a statistical technique used to reduce the dimensionality of data while retaining as much variance as possible. It works by transforming the original variables into a new set of uncorrelated variables called principal components (PC), which are ordered by the amount of variance they capture. The first few principal components often capture the majority of the data’s information, allowing for data compression and simplifying analysis. A comprehensive review of PCA’s applications and theory was published by Jolliffe.^56^ There are two principal components in our PCA study. The first component comprises the pairwise Euclidean distances between all C*_α_* atoms, yielding a total of (164 × 163)/2 features. These distances describe the relative spatial variations among internal atoms throughout the protein, thereby capturing global conformational differences. The second component is the distance between the centers of mass of residue groups 900-981 and 200-281, providing a single feature that reflects the conformational changes of the second alpha-helix (close to the N-terminal of MYC). Subsequently, this collective variable matrix is reduced to two principal components to facilitate visualization and clustering.

#### The R-package

Metadynminer was used to analyze the WTM results, such as calculating the free energy surface, finding basins of minimal free energy and analyzing transition pathways between stable states (minimum free energy paths, MFEP) in order to describe the most probable transition routes between these states on the FES and to eventually locate the transition states (TS) of the system.^57^ The MFEP represent the reaction pathways with the lowest energy demand between two stable states.

The *neb* function of metadynminer also estimates the half-life of these states ^58^ using the classical Eyring–Polanyi equation of chemical kinetics,^59^ with a transmission coefficient assumed to be 1. Hence, transition half-life is calculated from

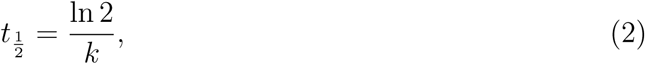

where *k* is derived from the Eyring–Polanyi equation. It represents how long a system remains in one stable state before transitioning across an activation barrier to another. In other words, it is the time required for half of a starting population of molecules to cross from a stable state through an energy barrier into a new stable state. Finally, the software VMD^60^ was used for visualization and analysis purposes.

## Results and discussion

This Section has been distributed into three main blocks. The first block is devoted to the construction of MD trajectories generated by starting from three different crystal structures obtained from PDB sources. The second one describes the two-dimensional free energy surface of MYC with full details, including the calculation of minimum free energy paths between stable states and the location of the most relevant transition states between stable states, together with its projection onto each CV. The third part is devoted to the characterization of the potentially targettable drugging pockets, revealed as stable basins located at the FES and we will close the ”Results” Section with a study of the docking of the main stable structures with a prototype small-molecule inhibitor and the simulation of the resulting complexes to validate the stability of the drugging pocket.

### Stable structures from Principal Component Analysis

MD was conducted on six parallel runs, yielding a total of 15 *µ*s of trajectory data. The initial structures of the three MYC motifs were derived from the crystal structures ’1NKP’, ’8OTS’ and ’6G6K’, where ’1NKP’ and ’8OTS’ represent the MYC-MAX-DNA complex, and ’6G6K’ is the MYC-MAX complex. In each case, the MYC structure was isolated from its partners and extracted. The results from MD simulations are summarized in Figure 3 and include the average of the three longest trajectories.

**Figure 3:**
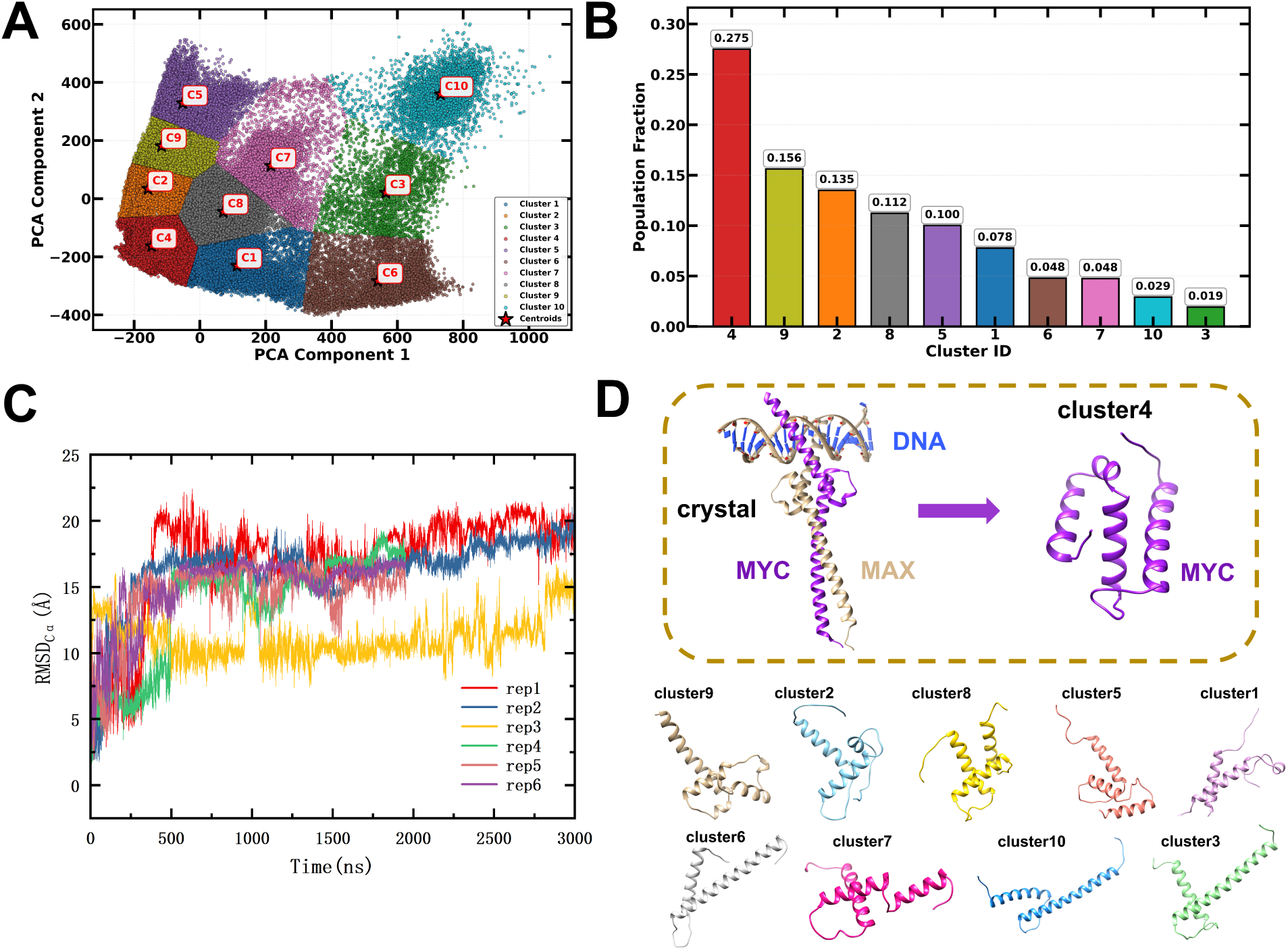
PCA clustering averaged from six independent MD runs, initialized with three different crystal structures ’1NKP’, ’6G6K’ and ’8OTS’. (A) Ten distinct clusters identified from PCA results, with the representative structure of each cluster (centroid) indicated by a red label; (B) Cluster density distributions; (C) RMSD profiles over the full simulation time of the six MD simulations, spanning a total of 3 *µ*s (replicas 1-3) or 2 *µ*s (replicas 4-6); (D) Most stables structures (clusters 1-10) ordered by cluster density (top to bottom, left to right).

We can observe (Fig.3, panels A-B) that structure ”4” is clearly the most populated of the ten detected ones, what indicates that such structure might correspond to the most stable one, holding for the largest part of the MD trajectories. Other structures such as those numbered as ”9”, ”2”, ”8” and ”5” have been recorded more than 10% of the total number of configurations analyzed. Remarkably, the five structures highlighted are qualitatively different of the CS of c-MYC published at RCSB PDB, more similar to structures ”6”, ”10” and ”3” also reported in Fig.3.

Regarding the high flexibility of the MYC monomer structure, it is essential to obtain sufficient and diverse conformations to identify relatively stable structures. For this purpose, our study employs WTM as a strategy to enhance sampling. This method assists MYC in overcoming energy barriers by applying bias forces, thereby enabling a broader exploration of the conformational space. This will allow us to determine whether the conformations identified by PCA, being totally different to the experimentally determined CS, correspond to thermodynamically stable or metastable states, since in general the most populated structures obtained from conventional unbiased MD trajectories will not necessarily coincide with the most stable conformations identified through enhanced-sampling methods such as WTM.

### Two-dimensional free energy landscapes of MYC

The two dimensional (2D) free energy landscape of MYC is represented in Fig. 4 for the two selected CV. We can observe that several main basins (such as A, B,…G) and their corresponding molecular conformations are well defined in all cases. We have only indicated the basins which are clearly seen from the FES, assuming that even with well converged runs the clearest definition of the FES would require a huge amount of statistics out of the standard available ranges. A detailed study of the convergence of the WTM study is reported in the ”Supporting Information” (SI), Figures 8-10.

**Figure 4:**
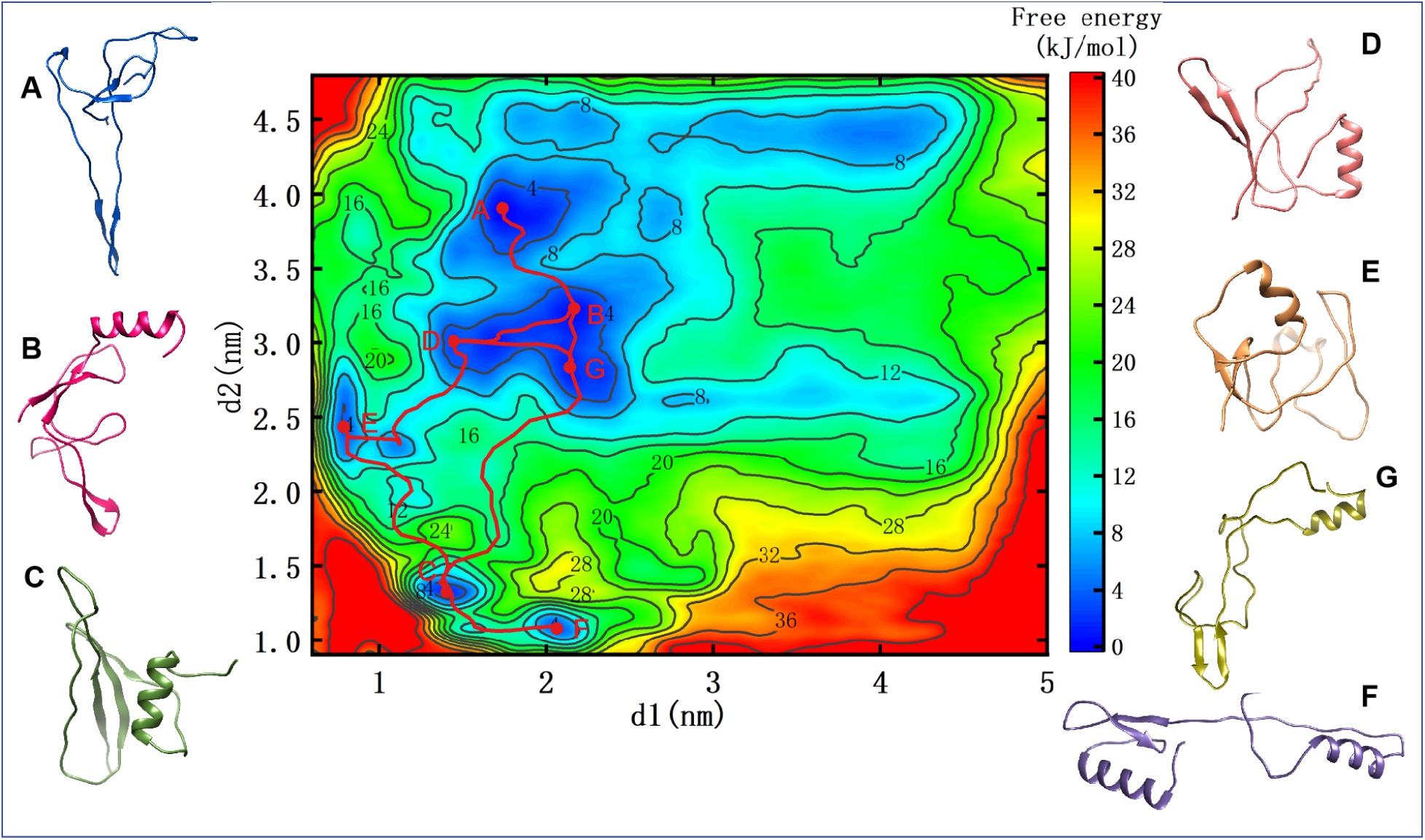
The 2D free energy landscape F(*d1*, *d2* ) (kJ/mol) for MYC in solution. Basins ”A”, ”B”, ”C”, ”D”, ”E”, ”F” and ”G” are the most thermodynamically stable states (minima) as they are explicitly indicated. The most visited conformations corresponding to the representative basins are listed on the two sides of the 2D free energy landscape. Minimum free energy paths are depicted in red lines.

**Figure 5:**
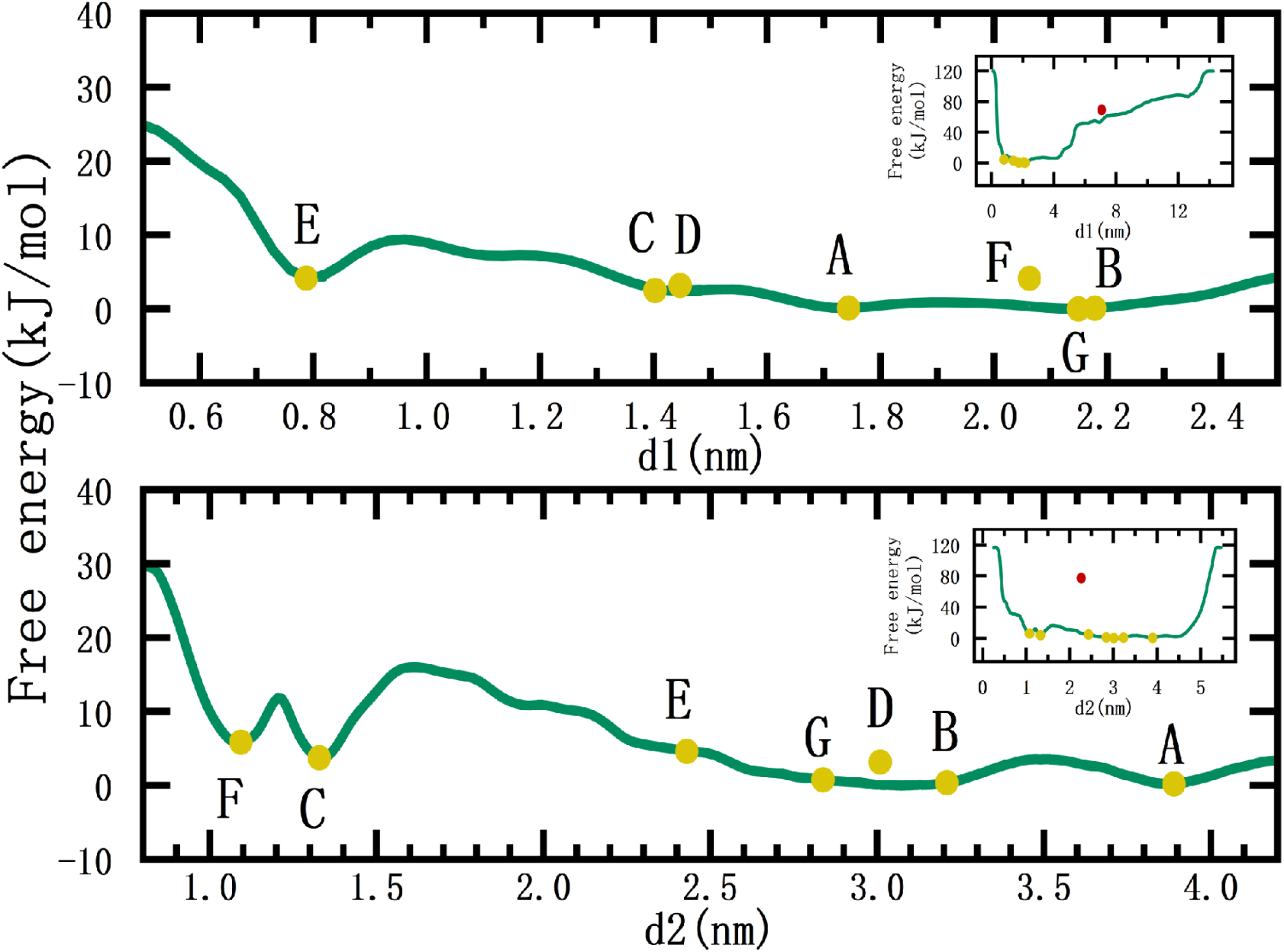
1D integrated free-energy profiles for two CV. Top row correspond to the distance CV *d*1, whereas bottom row correspond to the CV *d*2. Corresponding insets show the full range of values considered for each CV. Basin regions marked with same labels (A-G) as in Fig. 4. In order to directly compute the height of free-energy barriers, absolute minima are set equal to zero. The yellow dots indicate the location of the seven lowest energy minima, whereas the red dots indicate the location of the CS.

**Figure 6:**
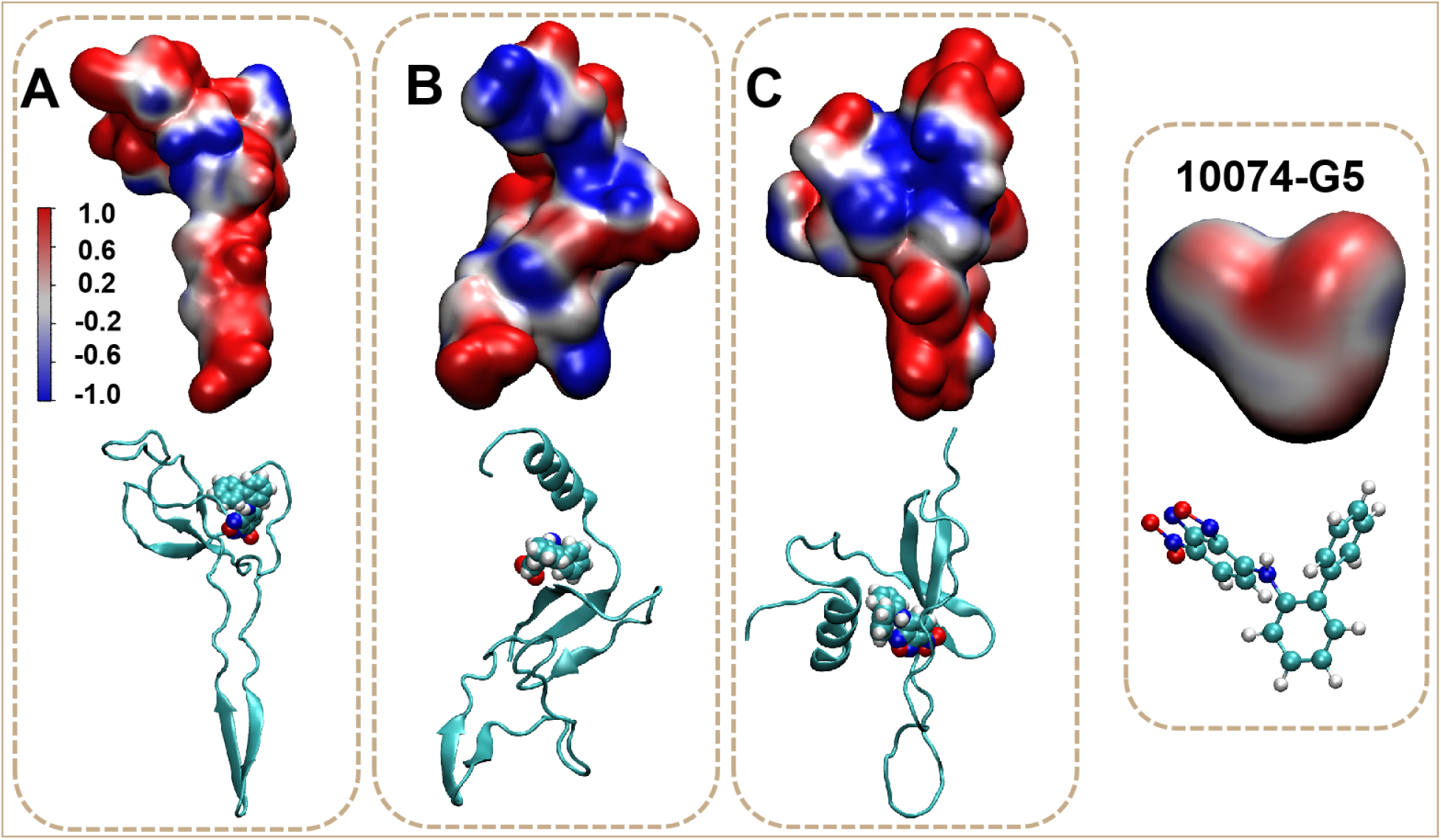
Electrostatic potential of selected MYC conformations in free energy basins A, B, and C (as reported in the 2D FES, Fig. 4) in units of *k_B_T/e*, where 1 *k_B_T/e* ≈ 26.7 mV, highlighting positive (red) and negative (blue) side chains of each pocket. The most suitable drug-binding pocket of each state will be tested by allocating the small-molecule inhibitor 10074-G5,^67^ whose charge distribution and atomic structure are shown at the right side of the figure (carbon atoms in cyan, hydrogens in white, nitrogens in dark blue and oxygens in red).

**Figure 7:**
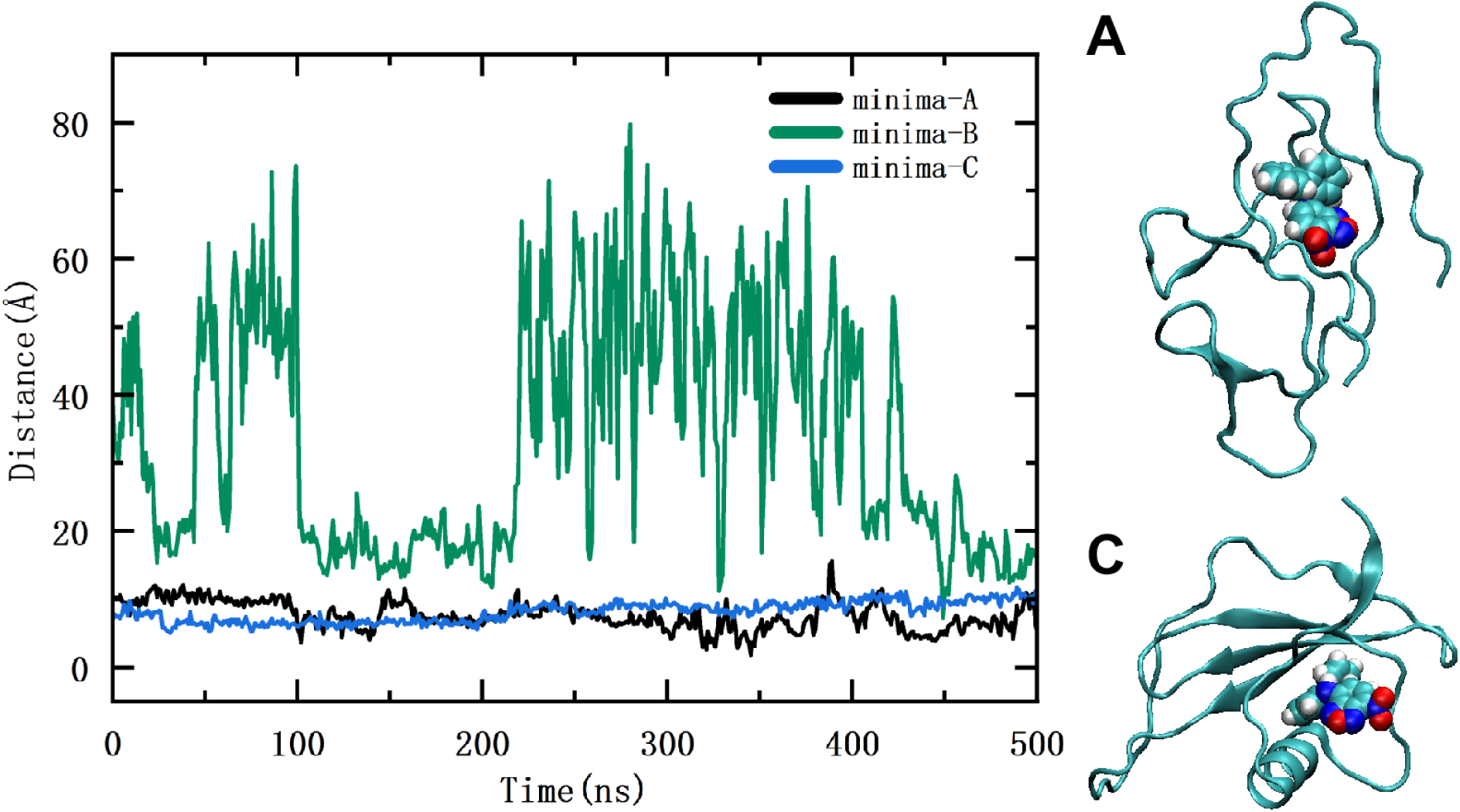
Time evolution of the distance between the center of mass of the MYC motif and the center of mass of the prototype drug 10074-G5 in the case of the three minima A, B, C (as indicated in Fig. 4). In the case of stable binding (configurations A, C) a snapshot of the final configuration is shown. The inhibitor 10074-G5 is displayed in van der Waals representation and all figures were generated using VMD.

**Figure 8:**
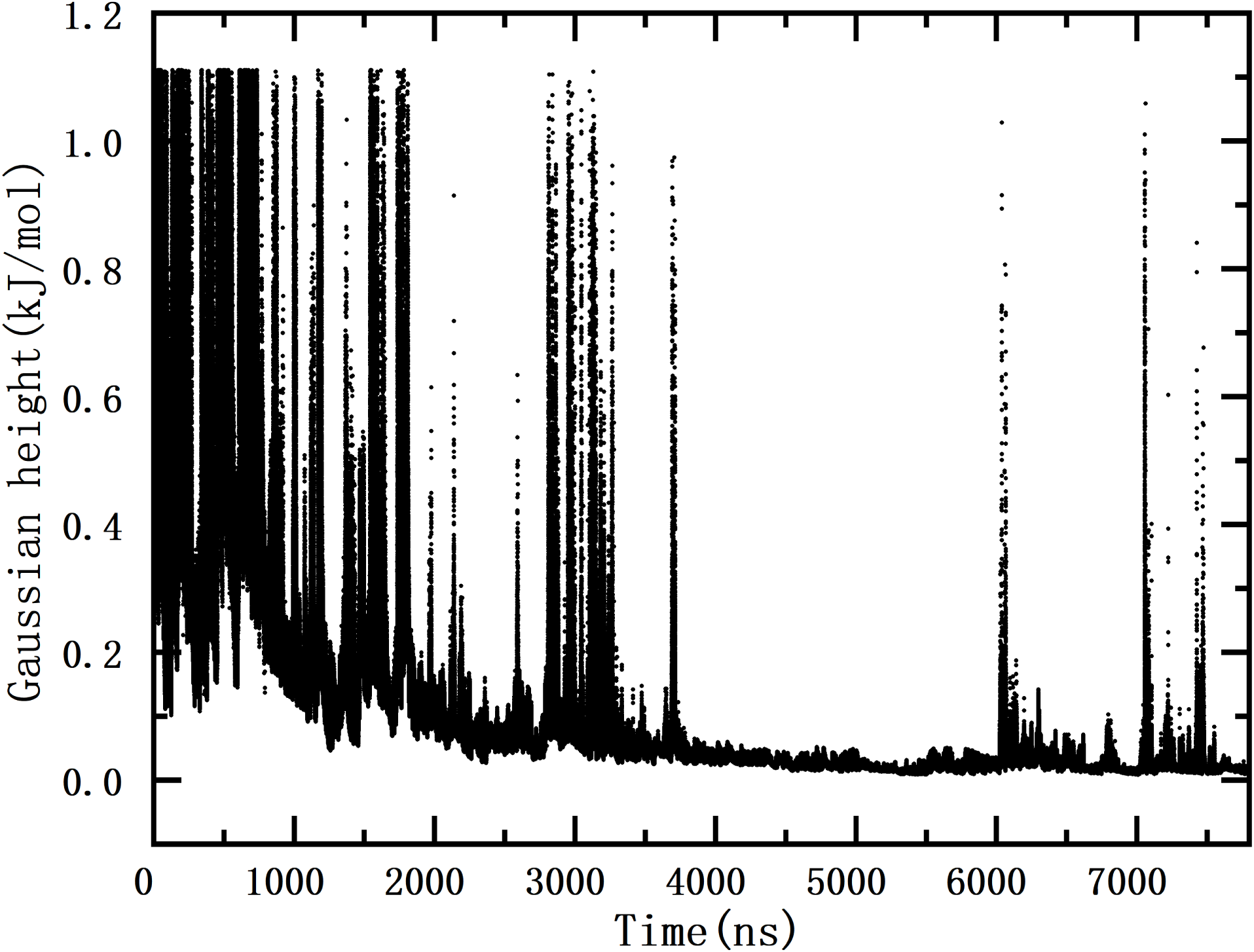
Time-evolution of WTM’s hills.

**Figure 9:**
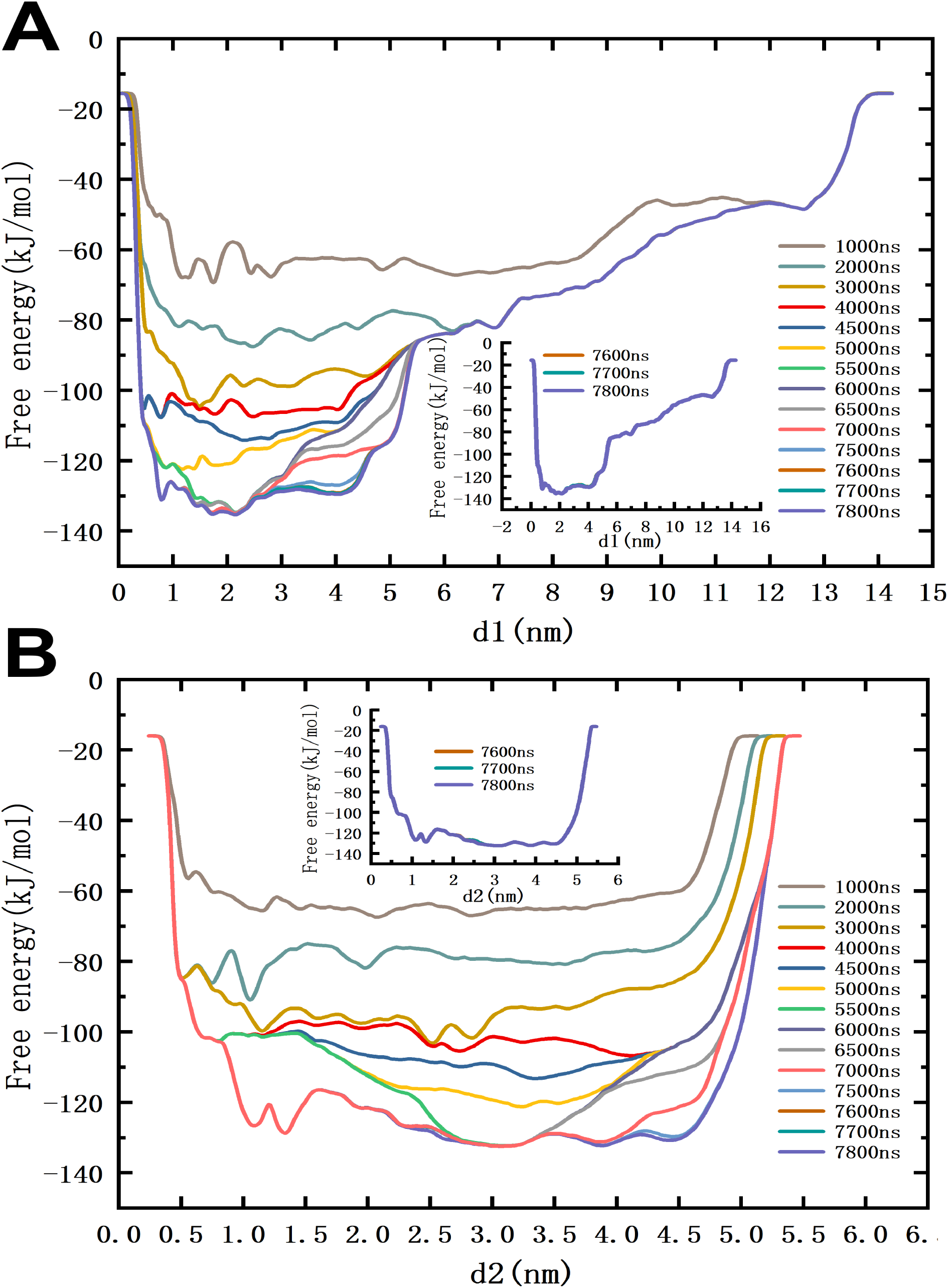
Convergence of one-dimensional free energies at different simulation times. (A) Free energy as a function of *d1* (B) free energy as a function of *d2*.

**Figure 10:**
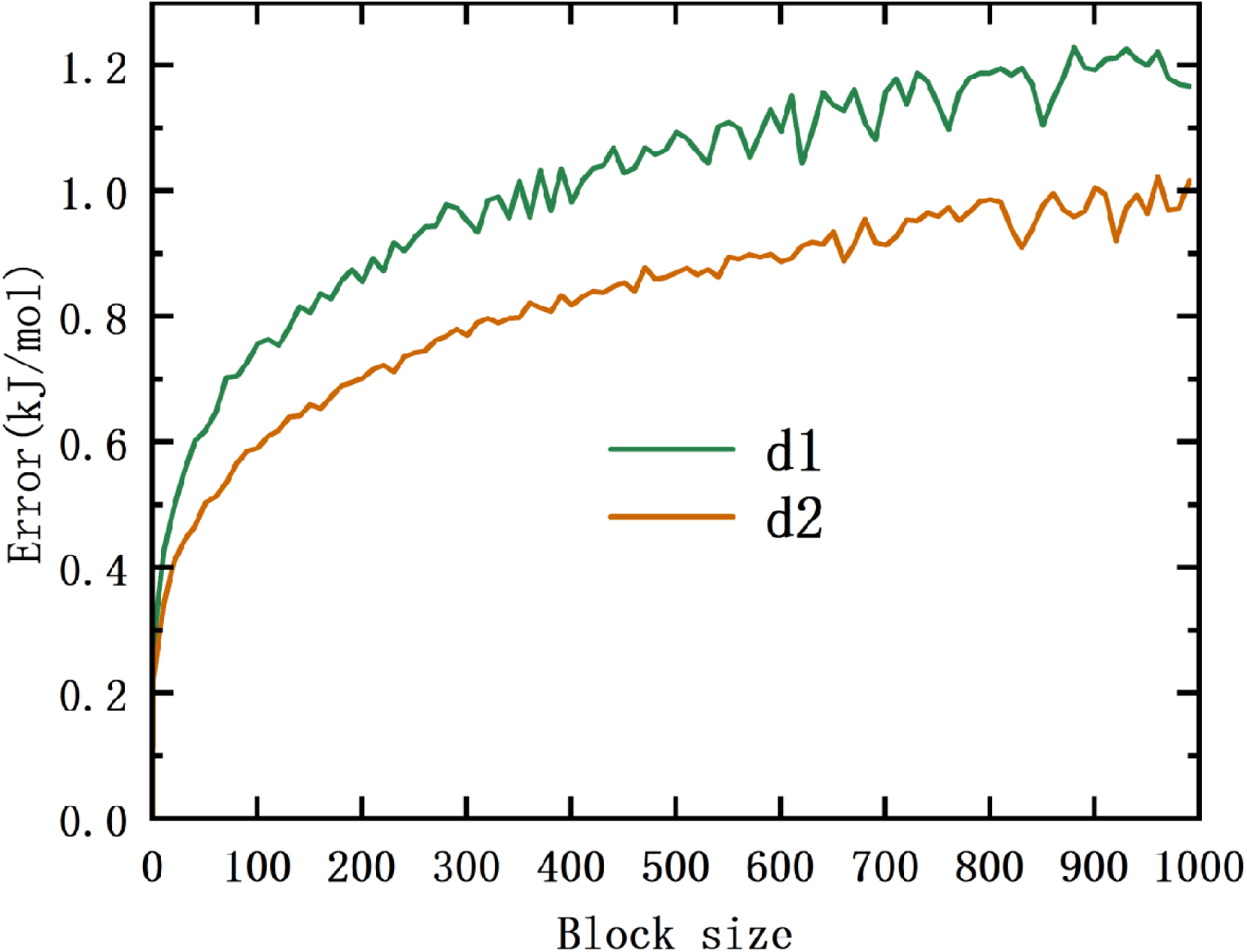
Block-size averages, for the two CV employed in this work.

As a general concept, the basins in the FES indicate the most probable configurations of the system in equilibrium. In fact, WTM is able to reveal those stable configurations in such a way that the free energy barriers that the system needs to overcome to shift between stable states can be estimated with a high degree of accuracy. In this work, we can distinguish several deep basins located around values of *d1* between 0.79 and 2.17 nm and distances *d2* between 1.08 and 3.90 nm. The coordinates of every selected basin (A-G) are reported in

Table 2, together with their free energy values, where the minimum of all them (state A) has been set to zero, producing the corresponding rescaling of the remaining ones.

**Table 2:** Coordinates and free energies of the representative basins (minima) corresponding to the Fig. 4, compared to the CS.

| Free energy [kJ/mol] | Basin | $d1$ [nm] | $d2$ [nm] |
| --- | --- | --- | --- |
| 0.00 | A | 1.74 | 3.90 |
| 1.01 | B | 2.17 | 3.23 |
| 2.49 | C | 1.40 | 1.33 |
| 3.18 | D | 1.45 | 3.01 |
| 3.93 | E | 0.79 | 2.44 |
| 4.14 | F | 2.06 | 1.08 |
| 1.33 | G | 2.13 | 2.88 |
| 74.69 | CS | 7.26 | 2.30 |

As a general fact, the lowest free energy basins show values of *d1* in a tight short range around 1-2 nm, indicating that in all cases the distance between C- and N-terminals of MYC is quite short, suggesting ”closed” and ”packed” structures. Further, the wider range for *d2* (about 1-4 nm) suggests a variety of different structures and shapes for the second alpha helix of MYC. Noticeably, the reported coordinate *d1* of the CS is much larger than the ones of the stable basins, whereas *d2* for CS falls right inside the same range of those of stable basins. This means that CS would fall in a high-energy region of the 2D FES (not reported in Fig. 4).

Table 2 shows that basins A, B, C and G correspond to the regions of lowest free energy (0-2.5 kJ/mol) and may represent stable states that are structurally related to the native ensemble, but including relevant differences in loop orientations or side-chain packing. Otherwise, basins D-F are located in slightly higher-energy regions (between 3-5 kJ/mol), corresponding to less stable conformations, although these are basins that could be considered as containing potential drugging pockets as well. States with free energies beyond 10 kJ/mol have not been considered. Furthermore, we observe (Table 2) a substantial free energy difference of ∼ 75 kJ/mol between the most stable basin (A) and the experimentally resolved CS, which may be of great relevance for the development of drugs targeting MYC. It is worth noticing that during the entire WTM simulation, MYC did not recover its initial level of helicity, even though regions of the free energy landscape corresponding to the same CV values were revisited multiple times, suggesting that the crystal structure may not represent the most probable configuration in that region. For such a reason, designing inhibitors that target these low-energy conformations is potentially more promising in order to achieve stable binding between the MYC protein and the inhibitor. Conversely, templating on the crystal structure is likely to result in failed binding due to MYC’s spontaneous transition to lower-energy states. This provides a clear thermodynamic rationale for selecting structures A–G as target conformations. In passing, we should notice that a triplet formed by sub-states B, D and G is formed by three states located in a small region with similar free energy values and short transition half-life.

### Identification of transition paths and transition states

Generally, basins in the FES indicate the most probable configurations of the system in equilibrium. As a matter of fact, WTM allows us to estimate the free energy barriers that the system needs to overcome to shift between stable states with a high degree of accuracy. Furthermore, maxima along each path give indications of the approximated location of TS between stable states. For instance, the TS between basins A and B is located around *d*_1_ ∼ 1.8 nm, *d*_2_ ∼ 3.6 nm and it is of about 7.2 kJ/mol. The full set of free energy profiles along the MFEP for each transition case are shown in the SI (Fig. 11), including the estimated locations of local TS between stable basins. Regarding the stability and readiness of MYC to transit between the located basins, we report in Table 3 the free energy barriers for the basins of interest located from Fig. 4, together with transition half-lives between these basins, estimated using the Eyring–Polanyi equation (see ”Methods”).

**Figure 11:**
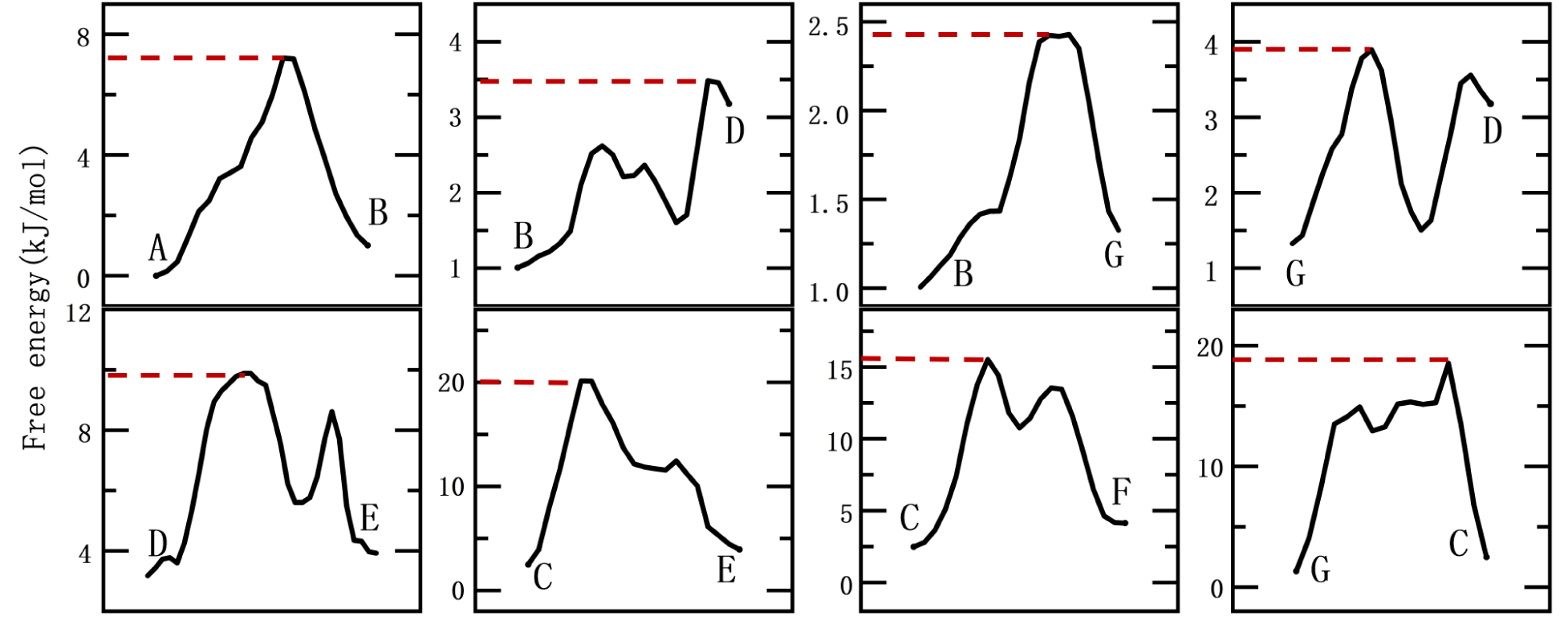
Free energy profiles evaluated over the minimum free energy paths shown in Fig. 4 of the main text. TS are indicated with a dashed line.

**Table 3:** Free energy barriers (ΔF) between basins (kJ/mol) along the corresponding MFEP and their transition half-lives (ps), as predicted by the Nudged Elastic Band method^61^ and Eyring–Polanyi equation, respectively (see Section ”Methods”). State transitions with half-lives shorter than 1 ps are shown in bold characters.

| State transition | $\Delta F$ (kJ/mol) | Transition half-life (ps) |
| --- | --- | --- |
| A to B | 7.21 | 1.76 |
| B to A | 6.21 | 1.19 |
| B to D | 2.48 | **0.28** |
| D to B | 0.31 | **0.12** |
| B to G | 1.42 | **0.19** |
| G to B | 1.10 | **0.16** |
| D to G | 0.72 | **0.14** |
| G to D | 2.57 | **0.29** |
| D to E | 6.70 | 1.45 |
| E to D | 4.69 | **0.66** |
| E to C | 16.21 | 57.77 |
| C to E | 17.65 | 101.02 |
| C to F | 13.04 | 16.92 |
| F to C | 11.40 | 8.93 |
| C to G | 13.01 | 16.68 |
| G to C | 8.69 | 3.12 |

The energy barriers between low-energy basins and higher-energy regions (e.g., D to E, C to E, C to F and G to C) suggest that some large-scale rearrangements are thermodynamically unfavorable and would require to cross high activation barriers. Conversely, energy barriers between the lowest energy basins (e.g., A to B, B to D, B to G and D to G) are within a moderately narrow range of a few kJ/mol and indicate that some rearrangements thermodynamically favorable would require to cross low activation barriers. Half-life values along the MFEP between basins are consistent with the number of conformers sampled for each basin in the WTM trajectories and short half-lifes show faster kinetic transitions between stable states. In particular, structures of MYC corresponding to basins exhibiting very short transition half-lives (less than 1 ps) indicate that they are transient states, i.e. resonances between two states of the same physical structure and meaning such as, for instance, interconversion flips between chemical isomers. We have detected the transitions between B, D and G, with half-lives lower than 0.3 ps, what allow us to categorize the triplet of these 3 substates as the same physical state. The largest barriers (mainly involving transitions between C and E, F or G) indicate strong stability of these states. State A shows a much lower barrier but it only can transit to state B, so that we can recall that states A, C and B-D-G are the best initial candidates for the search of drugging pockets on their surface.

Overall, the results for the free energy barriers reported here show many possible transitions between free energy basins. For instance, the transition half-lives between B, D and G are all less than 0.3 ps, with energy barriers lower than the thermal motion energy corresponding to 310 K (∼ 2.6 kJ/mol), indicating that they can rapidly interconvert under physiological conditions.

### One-dimensional free energy landscapes of MYC

The 2D free energy landscape obtained from WTM simulations (Fig. 4) has been build using the two CV defined above. Nevertheless, it is possible to obtain one-dimensional (1D) profiles where free energy depends of only one CV, after the second CV has been integrated out. This sort of calculations will allow us to directly compare free energy barriers to experimental findings (see Jämbeck et al. ^62^), since information about binding modes in experimental solutes is usually not available. Then, it is possible to use experimental binding free energies Δ*G_bind_* to get indirect leverage on the quality of the computed free energy barriers. In this work, 1D free-energy profiles F(*s*_1_) (comparable to the experimental Δ*G_bind_*) can be obtained by means of the following expression:^62,63^

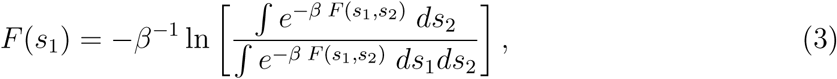

where *s*_1_ and *s*_2_ are the CV, *β* = 1*/*(*k*_B_*T* ), *k*_B_ is the Boltzmann constant and T is the absolute temperature. Here, the free energy *F* (*s*_1_) depends only of *s*_1_, whereas all possible contributions due to *s*_2_ have been integrated out and averaged. The results reported in Fig. 5 reveal a series of free energy barriers around the stable basins, most of them located very close to the lowest free energy state (A).

The two free energy profiles F(*d*_1_) and F(*d*_2_) are quite smooth, showing very close values for the stable states (A-G). Most basins can be reached by crossing very low free energy barriers, as indicated in Table 3. The full free energy landscape spans *d*1 from 0 to 14.3 nm and *d*2 from 0 to 5.5 nm (inseets in Fig. 5). States A-G are concentrated in the low energy region of coordinate *d*1, reflecting the higher stability of compact structures. In contrast, states A-G are more widely distributed along the coordinate *d*2, providing a basis for MYC conformational diversity. All selected structures, excepting states E and F, have reached their lowest free energy for *d*1 or *d*2. Nevertheless, the CS is located at *d*1 = 7.26 nm, *d*2 = 2.30 nm, very far away from the most stable conformations, exhibiting a large difference of approximately 75 kJ/mol. We should notice that values of *d*1 of states F, B and G overlap, with B and G exhibiting lower energies. This results in point F for *d*1 of Fig. 5 lying slightly above the free energy curve. A similar situation occurs for state D in the free energy projection on *d*2. Further, the energy barriers for F and C are primarily originated from the dependence on *d*2, with their free energy surface exhibiting a characteristic double-well. In summary, states A and B are the globally most stable structures (free energy difference of only 1.016 kJ/mol), whereas state C represents a moderately low free energy but structurally different of A and the triplet B-D-G.

Overall, the results for the free energy barriers reported here show many possible transitions between free energy basins involving free energies larger than the thermal energy at 310 K (∼ 2.6 kJ/mol). These barriers can be compared to other biological bottlenecks, such as those related to the RAS family of oncogenes, where barriers of ∼ 20 kJ/mol due to changes in specific distances and orientations were estimated for the activation of mutated RAS proteins by Fourier Transform Infrared Spectroscopy^64^ and simulations^53^ or to the case of KRAS-G12D, when bound to a cell membrane, where angular barriers in the range 8-17 kJ/mol were estimated.^65^

### Location of potential drugging pockets at the surface of MYC

A possible way to determine the location of potential drugging pockets at the surface of the stable structures of MYC previously reported in Sections ”Two-dimensional free energy landscapes of MYC” and ”One-dimensional free energy landscapes of MYC” is by plotting the charge densities at the MYC surface obtained from electrostatic calculations using the Poisson-Boltzmann equation. We have performed this calculation by means of the tool APBS (Adaptive Poisson-Boltzmann Solver^66^). The results are presented in Fig. 6 and indicate the existence of clear spots where charge is neatly positive (red) and others where it is mostly negative (blue), whereas neutral regions are plotted in white.

According to the selection of stable basins indicated above (namely minima A, B and C in Fig. 4), we hypothesize that these states should be considered by their potential as promising therapeutic targets in MYC-associated diseases. Minima A, B and C not only correspond to the lowest energy in the potential landscape but also represent three relatively independent states with distinct conformations. Among these, B stands for three interconvertible subconformations: B, D and G. As it can be seen in Fig. 6, there exist large pockets of these structures with electrostatic potential differences up to 2 *k_B_T/e*, i.e. ∼ 53 mV. These negatively charged regions are complementary to the overall positive charge exhibited by the small molecule 10074-G5, indicating that they can form electrostatic interactions. 10074-G5 (N-([1,1’-biphenyl]-2-yl)-7-nitrobenzo[c]oxadiazol-4-amine)^68^ is a MYC small-molecule inhibitor that impedes MYC-Max complex formation by binding to MYC, consequently attenuating the proliferative activity of tumor cells.^67^**^?^** In a similar fashion as it was described for MAX,^55^ we can assume that electrostatic interaction, say mediated by hydrogen bonds or salt bridges, might be a suitable mechanism for a small-molecule drug binding to MYC.

Our results suggest that the electrostatic binding of 10074-G5 to states A, B (resonance of triplet B-D-G), and C of MYC should be feasible given their different polar regions. In this way, we have conducted additional docking analyses between MYC and the 10074-G5 inhibitor, as predicted by Autodock Vina.**^?^ ^?^** The results of the docking analysis are reported in the SI (Table 4). The results, included in Fig. 6, show the most favorable complex structures for states A, B, and C and agree well with our preliminary predictions. In order to study the stability of the complexes formed by the three states bound to the 10074-G5 inhibitor, we have run three independent MD simulations and studied the distances between the center of mass of the 10074-G5 inhibitor and the center of mass of each MYC motif. The results of 500 *ns* runs are shown in Fig. 7. State B tends to exhibit significant fluctuations, reflecting transient departures of the inhibitor from the center of mass of MYC followed by occasional returns on short timescales. These observations suggest that state B corresponds to a thermodynamically stable basin while remaining dynamically flexible, consistent with the short transition half-life observed between basin B and neighboring basins. In comparison, the binding of the 10074-G5 inhibitor into the drug pockets in states A and C of MYC are very stable, with distances located around 1 nm during the full period of simulation time. This result suggests that, for the particular inhibitor 10074-G5, at least two drugging pockets reported in the present work are able to host this drug in a stable way during periods at the time scale of 500*ns*. As an additional test, we have considered other higher-energy minima such as E and F, together with the D and G (B’s partners) and found that their stability as drugging pockets is not very much different of that of A, B and C, when the inhibitor 10074-G5 is considered (see Fig. **??** in SI). Given such preliminary promising results, we are currently working in the computational design of new prototype inhibitors, that will be fully tested in the near future.

**Table 4:**
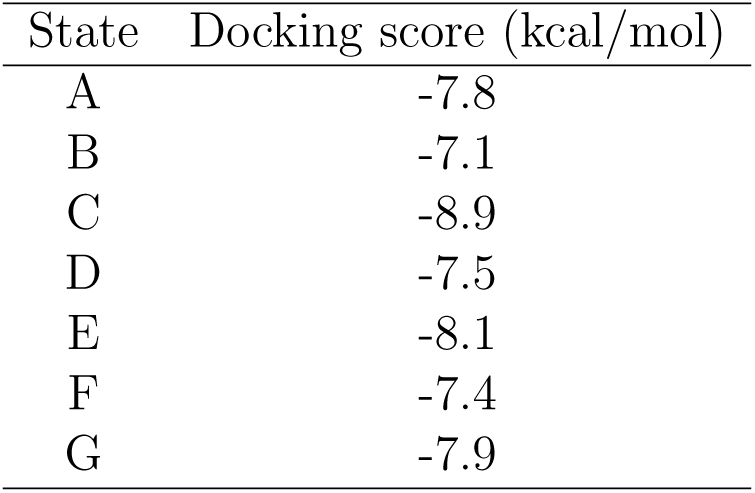
Docking scores.

| State | Docking score (kcal/mol) |
| --- | --- |
| A | -7.8 |
| B | -7.1 |
| C | -8.9 |
| D | -7.5 |
| E | -8.1 |
| F | -7.4 |
| G | -7.9 |

For MYC, which is typically regarded as ‘undruggable’ due to sequence disorder and the absence of stable drug pockets, this result holds significant importance: it not only validates the feasibility of minima A and C as potential targets but also provides a structural foundation for the subsequent design and screening of structure-based small-molecule inhibitors. Furthermore, It highlights the pivotal role of conformational dynamics in the evaluation of poorly druggable targets, and it paves the way to research into similar disordered proteins.

## Conclusions

This work reports a study of a motif c-MYC structures in solution. Determining the main conformations of this proto-oncogene is crucial, as it forms a MYC-MAX dimer that binds to DNA inside the human cell nucleus. This complex plays a key role in dysregulating gene expression in about 70% of all known cancers. Given the lack of clinical strategies directly targeting MYC, identifying specific druggable pockets on its surface or interior is highly valuable for drug design. Here we present a detailed study of the free energy surface of c-MYC in solution to locate previously uncharacterized druggable pockets. To achieve this, we have combined MD-WTM simulations with auxiliary tools, including minimum free energy path and docking analyses for a 10074-G5 inhibitor. We observed that the unbound MYC motif exhibits a rich conformational ensemble, revealed by extensive MD and WTM simulations. MD simulations revealed seven dominant structural clusters of monomer, very different from those in the initial crystal structures isolated from the complex MYC-MAX-DNA obtained by X-ray and electron microscopy. These structures guided the selection of two distance-based collective variables for WTM enhanced sampling. The resulting FES revealed a broad conformational space featuring numerous low-energy basins corresponding to (meta)stable states.

According to the findings reported in Fig. 4 and Fig. 5, our FES shows stable states with low free energy barriers located at three different regions, as mapped by the two CVs. The most stable basin (A) has a free energy of ∼ 74.69 kJ/mol below the experimentally resolved crystal structure, corroborating that the latter is a low-populated state in solution. The next two low-energy basins (B and C) have been also selected to explore potential drugging pockets. These stable conformations represent promising candidates for effective inhibitor targeting, moving beyond the long-studied MYC-MAX crystal structures. Finally, electrostatic potential maps of states A and C reveal polar pockets that showed good affinity and stability when tested against the small-molecule MYC inhibitor 10074-G5, highlighting potential targets for future drug development.

## Data and Software Availability

The raw data and analysis scripts that support the findings of this study can be found in the corresponding repository of Github: https://github.com/yanhong77634/myc. The repository contains input files for WTM simulations, coordinates of meta- and stable MYC structures (related to the use of the PLUMED plugin), scripts for PCA calculations, raw block analysis data, minimum free energy pathways between different basins, the optimized structure of 10074-G5, along with docking results for 10074-G5 and the most favorable MYC–inhibitor complex structures predicted through docking simulations.

## Acknowledgement

The authors thank financial support provided by the Spanish Ministry of Science, Innovation and Universities. This publication is a part of the I+D+i project with reference PID2024-157478NB-C32, founded by MCIN/AEI/10.13-039/501100011033 and “FEDER Una manera de hacer Europa”. Huixia Lu thanks the financial support by the ”Margarita Salas” grant which is funded by the European Union–NextGenerationEU. Yanhong Ge is a Ph.D. fellow from the China Scholarship Council (grant 202306230043). J.M. thanks the *Generalitat de Catalunya* for the support through the grant *Grup de Recerca SGR-Cat2021 Condensed, Complex and Quantum Matter Group* reference 2021SGR-01411 and to the Polytechnic University of Catalonia-Barcelona Tech through the funding AGRUPS. The authors thankfully acknowledge computer resources from MareNostrum5 supercomputer as well as technical support provided by BSC (RES-BCV-2024-2-0006, RES-BCV-2024-3-0013 and RES-BCV-2025-1-0008).

## Supporting Information Available

We will focus this complementary set of data on: (1) convergence of WTM simulations; (2) location of TS for each transition between basins; (3) list of docking scores and (4) stability of high-energy potential pockets.

## Convergence of WTD simulations

To ensure the reliability of the WTM simulations reported in the present work we can use several criteria. Three of the most used are: (1) the time evolution of the fluctuations of the hills of the biased potential along the full trajectory; (2) the time cumulative average of 1D free energy profiles and (3) the so-called block analysis of the average error along free energy profiles as a function of the block length.

The evolution of the hills is presented in Figure 8. We observe an overall decreasing behavior, eventually including high spikes during the transient part of the simulation (0-2000 ns). The height of the biased potential decreases accordingly along the simulation run, so that a quasi-flat profile is already seen around 4000 ns, including a few low-height spikes. Afterwards, a few high spikes are observed when the simulation reaches higher-energy regions, but such regions are of small importance and the fluctuations are not relevant. However, this is a clear but not yet conclusive indication of the convergence of the WTM runs.

Further, from the results of Figure 9 we can see that after long cumulative time lengths, the differences between a profile and the one immediately before are quite small (less than 20 kJ/mol), leading us to fully converged free energies for the two CV considered in the present study.

Finally, for the sake of reaching full safety in the convergence tests, we report the size of the average error in the free energy profiles calculated from WTM simulations as a function of the block size in Fig. 10. As expected, the errors increase with the block length until they reach a plateau for the two CVs. The average errors tend to stabilize around 1.2 kJ/mol for *d1* and 1 kJ/mol for *d2*, indicating consistent behavior over extended simulation times. Both criteria indicate that the WTM simulations performed in this study have converged. For this calculation, scripts provided by the PLUMED project^47,48^ have been employed.

### Free energy profiles along the Minimum Free Energy Paths

In the main text, we reported the minimum free energy paths connecting different free energy basins as identified by the Nudged Elastic Band method^61^ (Fig. 4). Although the free energy values can be identified from that figure, in order to facilitate the discussion, we show here the free energy profiles explicitly evaluated over these paths (Fig. 11). We indicate the TS found along these paths.

### 10074-G5 and MYC docking simulations

We used AutoDock Vina**^?^ ^?^** to perform docking simulations for basins A–G with the small-molecule inhibitor 10074-G5, in order to predict their binding modes and affinities. The results of the docking scores between MYC and the 10074-G5 inhibitor are listed in Table 4. AutoDock Vina was employed to dock 10074-G5 with these pockets. The complex structure with the best docking score is shown in Figure 6. The inherent scoring function of AutoDock Vina yields the predicted binding free energy, evaluating the binding affinity between ligand and target. Decreasing values indicate stronger predicted binding affinity. Guided by established thresholds in drug screening, molecules with scores below −7.0 kcal/mol are generally considered to possess strong binding potential. States A-G all qualified under this criterion.

### Stability of high-energy potential drugging pockets

In a similar fashion as the results presented in Fig. 6, we report here (Fig. 12) a stability study of pockets D, E, F, and G showing a bad stability performance overall.

**Figure 12:**
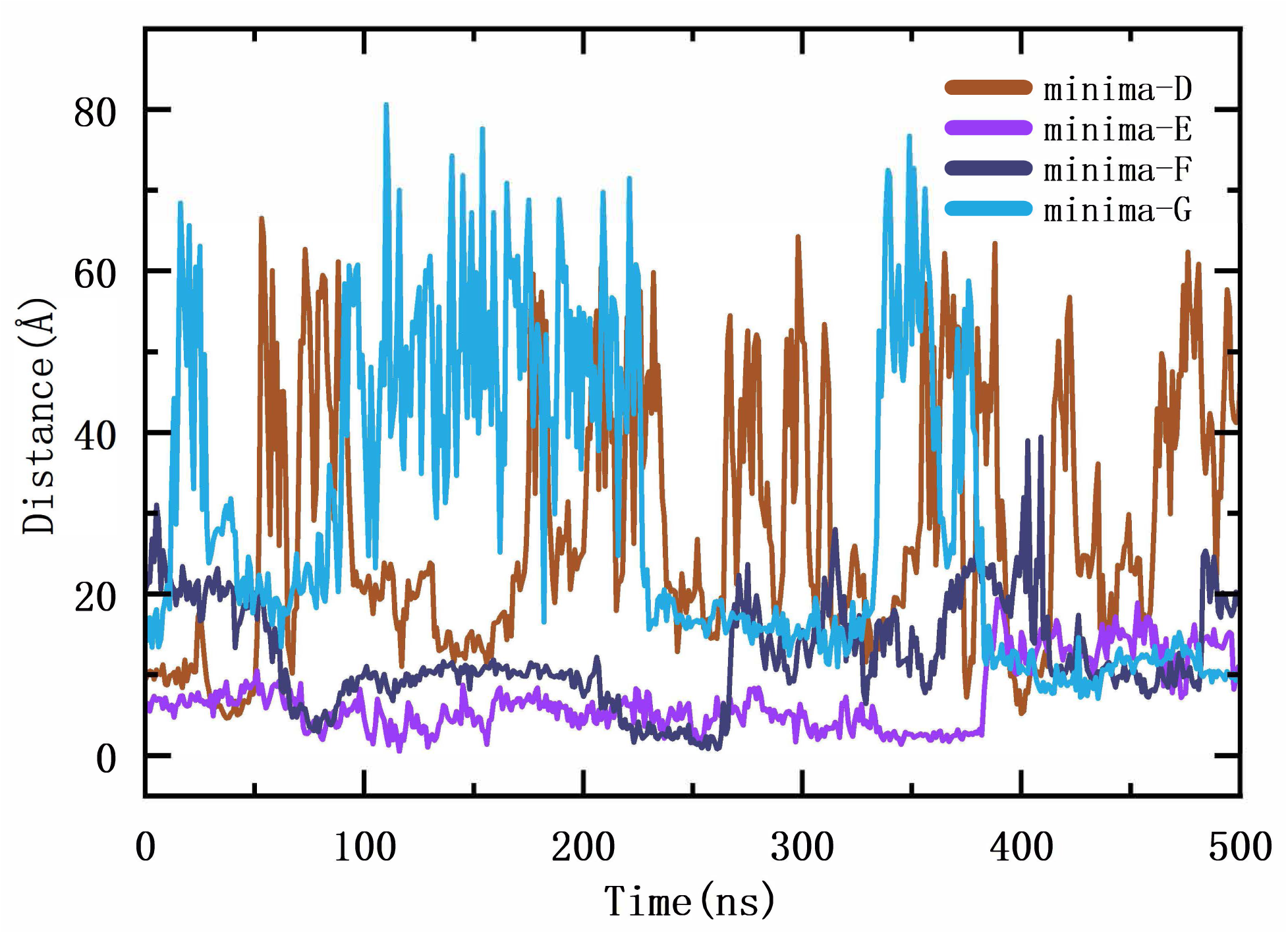
Time evolution of the distance between the center of mass of the MYC motif and the center of mass of the prototype drug 10074-G5 in the case of the minima D, E, F and G (as indicated in Fig. 4).

## Notes

### Competing Interest Statement

The authors have declared no competing interest.

## References

(1) Dang, C. V. MYC on the path to cancer. Cell 2012, 149, 22–35.

(2) Kortlever, R. M.; Sodir, N. M.; Wilson, C. H.; Burkhart, D. L.; Pellegrinet, L.; Swigart, L. B.; Littlewood, T. D.; Evan, G. I. Myc cooperates with Ras by programming inflammation and immune suppression. Cell 2017, 171, 1301–1315.

(3) Beaulieu, M.-E.; Castillo, F.; Soucek, L. Structural and biophysical insights into the function of the intrinsically disordered Myc oncoprotein. Cells 2020, 9, 1038.

(4) Duffy, M. J.; O’Grady, S.; Tang, M.; Crown, J. MYC as a target for cancer treatment. Cancer treatment reviews 2021, 94, 102154.

(5) Varmus, H. E. The molecular genetics of cellular oncogenes. Annual review of genetics 1984, 18, 553–612.

(6) Cole, M. D. The myc oncogene: its role in transformation and differentiation. Annual review of genetics 1986, 20, 361–384.

(7) Hermeking, H. The MYC oncogene as a cancer drug target. Current cancer drug targets 2003, 3, 163–175.

(8) Jakobsen, S. T.; Siersbæk, R. Transcriptional regulation by MYC: an emerging new model. Oncogene 2024, 1–7.

(9) Blackwood, E. M.; Eisenman, R. N. Max: a helix-loop-helix zipper protein that forms a sequence-specific DNA-binding complex with Myc. Science 1991, 251, 1211–1217.

(10) Atchley, W. R.; Fitch, W. M. Myc and Max: molecular evolution of a family of proto-oncogene products and their dimerization partner. Proceedings of the National Academy of Sciences 1995, 92, 10217–10221.

(11) Cascón, A.; Robledo, M. MAX and MYC: a heritable breakup. Cancer research 2012, 72, 3119–3124.

(12) Lourenco, C.; Resetca, D.; Redel, C.; Lin, P.; MacDonald, A. S.; Ciaccio, R.; Kenney, T. M.; Wei, Y.; Andrews, D. W.; Sunnerhagen, M.; others MYC protein interactors in gene transcription and cancer. Nature Reviews Cancer 2021, 21, 579–591.

(13) Ross, J.; Miron, C. E.; Plescia, J.; Laplante, P.; McBride, K.; Moitessier, N.; Möröy, T. Targeting MYC: From understanding its biology to drug discovery. European Journal of Medicinal Chemistry 2021, 213, 113137.

(14) Llombart, V.; Mansour, M. R. Therapeutic targeting of “undruggable” MYC. EBioMedicine 2022, 75 .

(15) Li, Z.; Huang, Y.; Hung, T. I.; Sun, J.; Aispuro, D.; Chen, B.; Guevara, N.; Ji, F.; Cong, X.; Zhu, L.; others MYC-targeting inhibitors generated from a stereodiversified bicyclic peptide library. Journal of the American Chemical Society 2024, 146, 1356–1363.

(16) Whitfield, J. R.; Soucek, L. MYC in cancer: from undruggable target to clinical trials. Nature Reviews Drug Discovery 2025, 1–13.

(17) Kiessling, A.; Sperl, B.; Hollis, A.; Eick, D.; Berg, T. Selective inhibition of c-Myc/Max dimerization and DNA binding by small molecules. Chemistry & biology 2006, 13, 745–751.

(18) Wang, C.; Zhang, J.; Yin, J.; Gan, Y.; Xu, S.; Gu, Y.; Huang, W. Alternative approaches to target Myc for cancer treatment. Signal transduction and targeted therapy 2021, 6, 117.

(19) Kota, J.; Chivukula, R. R.; O’Donnell, K. A.; Wentzel, E. A.; Montgomery, C. L.; Hwang, H.-W.; Chang, T.-C.; Vivekanandan, P.; Torbenson, M.; Clark, K. R.; others Therapeutic microRNA delivery suppresses tumorigenesis in a murine liver cancer model. Cell 2009, 137, 1005–1017.

(20) Soucek, L.; Jucker, R.; Panacchia, L.; Ricordy, R.; Tato, F.; Nasi, S. Omomyc, a potential Myc dominant negative, enhances Myc-induced apoptosis. Cancer research 2002, 62, 3507–3510.

(21) Padró, J.; Marti, J.; Guardia, E. Molecular dynamics simulation of liquid water at 523 K. Journal of Physics: Condensed Matter 1994, 6, 2283.

(22) Padro, J.; Marti, J. Response to “Comment on ‘An interpretation of the low-frequency spectrum of liquid water’”[J. Chem. Phys. 118, 452 (2003)]. The Journal of Chemical Physics 2004, 120, 1659–1660.

(23) Calero, C.; Marti, J.; Guàrdia, E. 1H nuclear spin relaxation of liquid water from molecular dynamics simulations. The Journal of Physical Chemistry B 2015, 119, 1966–1973.

(24) https://www.uniprot.org/uniprotkb/P01106/entry#sequences.

(25) Deshpande, N.; Addess, K. J.; Bluhm, W. F.; Merino-Ott, J. C.; Townsend-Merino, W.; Zhang, Q.; Knezevich, C.; Xie, L.; Chen, L.; Feng, Z.; Kramer Green, R.; Flippen-Anderson, J. L.; Westbrook, J.; Berman, H. M.; Bourne, P. E. The RCSB Protein Data Bank: a redesigned query system and relational database based on the mmCIF schema. Nucleic Acids Research 2005, 33, D233–D237.

(26) Kouranov, A.; Xie, L.; de la Cruz, J.; Chen, L.; Westbrook, J.; Bourne, P. E.; Berman, H. M. The RCSB PDB information portal for structural genomics. Nucleic acids research 2006, 34, D302–D305.

(27) Jorgensen, W. L.; Chandrasekhar, J.; Madura, J. D.; Impey, R. W.; Klein, M. L. Comparison of simple potential functions for simulating liquid water. The Journal of chemical physics 1983, 79, 926–935.

(28) Jo, S.; Kim, T.; Iyer, V. G.; Im, W. CHARMM-GUI: a web-based graphical user interface for CHARMM. Journal of computational chemistry 2008, 29, 1859–1865.

(29) Brooks, B. R.; Brooks III, C. L.; Mackerell Jr, A. D.; Nilsson, L.; Petrella, R. J.; Roux, B.; Won, Y.; Archontis, G.; Bartels, C.; Boresch, S.; others CHARMM: the biomolecular simulation program. Journal of computational chemistry 2009, 30, 1545–1614.

(30) Lee, J.; Cheng, X.; Jo, S.; MacKerell, A. D.; Klauda, J. B.; Im, W. CHARMM-GUI input generator for NAMD, GROMACS, AMBER, OpenMM, and CHARMM/OpenMM simulations using the CHARMM36 additive force field. Biophysical journal 2016, 110, 641a.

(31) Huang, J.; MacKerell Jr, A. D. CHARMM36 all-atom additive protein force field: Validation based on comparison to NMR data. Journal of computational chemistry 2013, 34, 2135–2145.

(32) Berendsen, H. J.; van der Spoel, D.; van Drunen, R. GROMACS: A message-passing parallel molecular dynamics implementation. Computer physics communications 1995, 91, 43–56.

(33) Senn, H. M.; Thiel, W. QM/MM methods for biological systems. Atomistic approaches in modern biology: from quantum chemistry to molecular simulations 2007, 173–290.

(34) Geissler, P. L.; Dellago, C.; Chandler, D.; Hutter, J.; Parrinello, M. Autoionization in liquid water. Science 2001, 291, 2121–2124.

(35) Dellago, C.; Bolhuis, P. G.; Geissler, P. L. Transition path sampling. Advances in chemical physics 2002, 123, 1–78.

(36) Hénin, J.; Fiorin, G.; Chipot, C.; Klein, M. L. Exploring multidimensional free energy landscapes using time-dependent biases on collective variables. Journal of chemical theory and computation 2010, 6, 35–47.

(37) Bartels, C.; Karplus, M. Multidimensional adaptive umbrella sampling: Applications to main chain and side chain peptide conformations. Journal of Computational Chemistry 1997, 18, 1450–1462.

(38) Barducci, A.; Bonomi, M.; Parrinello, M. Metadynamics. Wiley Interdisciplinary Reviews: Computational Molecular Science 2011, 1, 826–843.

(39) Bussi, G.; Gervasio, F. L.; Laio, A.; Parrinello, M. Free-energy landscape for *β* hairpin folding from combined parallel tempering and metadynamics. Journal of the American Chemical Society 2006, 128, 13435–13441.

(40) Deighan, M.; Bonomi, M.; Pfaendtner, J. Efficient simulation of explicitly solvated proteins in the well-tempered ensemble. Journal of chemical theory and computation 2012, 8, 2189–2192.

(41) Palmer, J. C.; Car, R.; Debenedetti, P. G. The liquid–liquid transition in supercooled ST2 water: a comparison between umbrella sampling and well-tempered metadynamics. Faraday discussions 2013, 167, 77–94.

(42) Haldar, S.; Kuhrova, P.; Banas, P.; Spiwok, V.; Sponer, J.; Hobza, P.; Otyepka, M. Insights into stability and folding of GNRA and UNCG tetraloops revealed by microsecond molecular dynamics and well-tempered metadynamics. Journal of Chemical Theory and Computation 2015, 11, 3866–3877.

(43) Martí, J. Free-energy surfaces of ionic adsorption in cholesterol-free and cholesterol-rich phospholipid membranes. Molecular Simulation 2018, 44, 1136–1146.

(44) Lu, H.; Marti, J. Cellular absorption of small molecules: free energy landscapes of melatonin binding at phospholipid membranes. Scientific reports 2020, 10, 9235.

(45) Lu, H.; Marti, J. Long-lasting Salt Bridges Provide the Anchoring Mechanism of Oncogenic Kirsten Rat Sarcoma Proteins at Cell Membranes. The Journal of Physical Chemistry Letters 2020, 11, 9938–9945.

(46) Martí, J.; Lu, H. Microscopic interactions of melatonin, serotonin and tryptophan with zwitterionic phospholipid membranes. International journal of molecular sciences 2021, 22, 2842.

(47) Bonomi, M.; Branduardi, D.; Bussi, G.; Camilloni, C.; Provasi, D.; Raiteri, P.; Donadio, D.; Marinelli, F.; Pietrucci, F.; Broglia, R. A.; others PLUMED: A portable plugin for free-energy calculations with molecular dynamics. Computer Physics Communications 2009, 180, 1961–1972.

(48) Tribello, G. A.; Bonomi, M.; Branduardi, D.; Camilloni, C.; Bussi, G. PLUMED 2: New feathers for an old bird. Computer physics communications 2014, 185, 604–613.

(49) Bussi, G.; Laio, A.; Parrinello, M. Equilibrium free energies from nonequilibrium meta-dynamics. Physical review letters 2006, 96, 090601.

(50) Barducci, A.; Bussi, G.; Parrinello, M. Well-tempered metadynamics: a smoothly converging and tunable free-energy method. Physical review letters 2008, 100, 020603.

(51) Bonomi, M.; Camilloni, C.; Vendruscolo, M. Metadynamic metainference: Enhanced sampling of the metainference ensemble using metadynamics. Scientific reports 2016, 6, 31232.

(52) Bussi, G.; Branduardi, D.; others Free-energy calculations with metadynamics: Theory and practice. Rev. Comput. Chem 2015, 28, 1–49.

(53) Hu, Z.; Martí, J. Atomic-level mechanisms of abnormal activation in NRAS oncogenes from two-dimensional free energy landscapes. Nanoscale 2025, 17, 4047–4057.

(54) Lu, H.; Marti, J. Influence of cholesterol on the orientation of the farnesylated GTP-bound KRas-4B binding with anionic model membranes. Membranes 2020, 10, 364.

(55) Lu, H.; Marti, J.; Faraudo, J. Probing the Structural Dynamics of the Unbound MAX Protein: Insights from Well-Tempered Metadynamics. Journal of chemical information and modeling 2025,

(56) Jolliffe, I. T.; Cadima, J. Principal component analysis: a review and recent developments. *Philosophical Transactions of the Royal Society A: Mathematical*, Physical and Engineering Sciences 2016, 374, 20150202.

(57) Hošek, P.; Spiwok, V. Metadyn View: Fast web-based viewer of free energy surfaces calculated by metadynamics. Computer Physics Communications 2016, 198, 222–229.

(58) Trapl, D.; Spiwok, V. Analysis of the Results of Metadynamics Simulations by metadynminer and metadynminer3d. *arXiv preprint arXiv:2009.02241* 2020,

(59) Eyring, H. The Activated Complex in Chemical Reactions. The Journal of Chemical Physics 1935, 3, 107–115.

(60) Humphrey, W.; Dalke, A.; Schulten, K. VMD: visual molecular dynamics. Journal of molecular graphics 1996, 14, 33–38.

(61) Henkelman, G.; Jónsson, H. Improved tangent estimate in the nudged elastic band method for finding minimum energy paths and saddle points. The Journal of chemical physics 2000, 113, 9978–9985.

(62) Jambeck, J. P.; Lyubartsev, A. P. Exploring the free energy landscape of solutes embedded in lipid bilayers. The Journal of Physical Chemistry Letters 2013, 4, 1781–1787.

(63) Roux, B. Statistical mechanical equilibrium theory of selective ion channels. Biophysical journal 1999, 77, 139–153.

(64) Li, Y.; Zhang, Y.; Großeruschkamp, F.; Stephan, S.; Cui, Q.; Kotting, C.; Xia, F.; Gerwert, K. Specific substates of Ras to interact with GAPs and effectors: revealed by theoretical simulations and FTIR experiments. The journal of physical chemistry letters 2018, 9, 1312–1317.

(65) Lu, H.; Marti, J. Predicting the conformational variability of oncogenic GTP-bound G12D mutated KRas-4B proteins at zwitterionic model cell membranes. Nanoscale 2022, 14, 3148–3158.

(66) Jurrus, E.; Engel, D.; Star, K.; Monson, K.; Brandi, J.; Felberg, L. E.; Brookes, D. H.; Wilson, L.; Chen, J.; Liles, K.; others Improvements to the APBS biomolecular solvation software suite. Protein science 2018, 27, 112–128.

(67) Yin, X.; Giap, C.; Lazo, J. S.; Prochownik, E. V. Low molecular weight inhibitors of Myc–Max interaction and function. Oncogene 2003, 22, 6151–6159.

(68) Madden, S. K.; de Araujo, A. D.; Gerhardt, M.; Fairlie, D. P.; Mason, J. M. Taking the Myc out of cancer: toward therapeutic strategies to directly inhibit c-Myc. Molecular Cancer 2021, 20, 3.

